# *De-novo* Genome Assembly of the Edwardsiid Anthozoan *Edwardsia elegans*

**DOI:** 10.1101/2024.10.02.616324

**Authors:** Auston Rutlekowski, Vengamanaidu Modepalli, Remi Ketchum, Yehu Moran, Adam Reitzel

**Affiliations:** Department of Biological Science, University of North Caroline at Charlotte, Charlotte, NC 28223; Center for Computational Intelligence to Predict Health and Environmental Risks, University of North Carolina at Charlotte, Charlotte, NC 28223; Marine Biological Association of the UK, The Laboratory, Citadel Hill, Plymouth PL1 2PB, UK; Whitney Laboratory for Marine Bioscience, University of Florida, St Augustine, FL 32080; Department of Genetics, University of North Carolina, Chapel Hill, North Carolina, USA; Department of Ecology, Evolution and Behavior, Alexander Silberman Institute of Life Sciences, Faculty of Science, The Hebrew University of Jerusalem, 9190401 Jerusalem, Israel

**Keywords:** Cnidaria, genome sequencing, synteny, Anthozoa, microRNA, genomic diversity

## Abstract

Cnidarians (sea anemones, corals, hydroids, and jellyfish) are a key outgroup for comparisons with bilaterial animals to trace the evolution of genomic complexity and diversity within the animal kingdom, as they separated from most other animals 100s of millions of years ago. Cnidarians have extensive diversity, yet the paucity of genomic resources limits our ability to compare genomic variation between cnidarian clades and species. Here we report the genome for *Edwardsia elegans*, a sea anemone in the most specious genus of the family Edwardsiidae, a phylogenetically important family of sea anemones that contains the model anemone *Nematostella vectensis*. The *E. elegans* genome is 396 Mb and predicted to encode approximately 49,000 proteins. We annotated large conservation of macrosynteny between *E. elegans* and other Edwardsiidae anemones as well as conservation of both microRNAs and ultra conserved noncoding elements previously reported in other cnidarians species. We also highlight microsyntenic variation of clustered developmental genes and ancient gene clusters that vary between species of sea anemones, despite previous research showing conservation between cnidarians and bilaterians. Overall, our analysis of the *E. elegans* genome highlights the importance of using multiple species to represent a taxonomic group for genomic comparisons, where genomic variation can be missed for large and diverse clades.

## Introduction

Species in the phylum Cnidaria have proven to be useful for understanding the evolutions of animals and their genomes (Putnam *et al*. 2007; Stefanik *et al*. 2014; Zimmermann *et al*. 2023). The first released cnidarian genome from the Edwardsiid sea anemone *Nematostella vectensis* revealed unexpected complexity in gene content and structure (Putnam *et al*. 2007), providing novel insights into the cnidarian-bilaterian ancestor. Since then, most reported genomes generated for the cnidarian phylum have focused on anthozoans (104 of 157 genomes), where the large majority (n = 94) are from Hexacorallia (stony corals and sea anemones). Within the Hexacorallia, only 20 of these species are sea anemones (Actinarians). The relative paucity of genomic data compared to the taxonomic and evolutionary diversity of sea anemones as well as the phylum Cnidaria more broadly, limits our understanding of genomic evolution in this ancient group and how it relates to the diversity of life histories represented by the many species of cnidarians. The lack of genomic data for much of the phylum leaves the community with the need to strategically sequence genomes from additional species to better resolve patterns and processes in genome evolution.

The family Edwardsiidae (Cnidaria, Anthozoan, Actinaria) contains over 100 species of sea anemones in 13 accepted genera, many composed of only a few species (World Register of Marine Species, accessed 04/09/2024). Species in the family Edwardsiidae are distributed throughout the world from polar to tropical seas, ranging from deep seas to shallow coastal habitats (Daly 2002; Mcfadden *et al*. 2007; Daly *et al*. 2008; Daly AND Ljubenkov 2008; Daly *et al*. 2013; Mcfadden *et al*. 2021). Edwardsiids have been studied due to unique structures (nemathybomes and nematosomes), their ‘simple’ anatomy and ‘phylogenetic primitiveness’ as adults, and challenges with resolving relationships of this family within Anthozoa (Daly 2002; Daly *et al*. 2002; Daly *et al*. 2008). Species of interest have included *Scolanthus callimorphus* (Zimmermann *et al*. 2023), the parasitic *Edwardsiella lineata* (Bumann AND Puls 1996; Stefanik *et al*. 2014), the Antarctic ice dwelling species *Edwardsiella andrillae* (Daly *et al*. 2013), and the model species for evolutionary developmental biology and genomics *Nematostella vectensis* (Darling *et al*. 2005; Layden *et al*. 2016; Al-Shaer *et al*. 2021). The most specious genus in this family is *Edwardsia* with more than 60 recognized species, which lacks any genomic data, limiting our ability to contextualize the genomic diversity and variation reported for existing genomes for *N. vectensis* and the recently published *S. callimorphus*. The genus *Edwardsia* diverged from other genera in the family 100s of millions of years ago (Mcfadden *et al*. 2021; Zimmermann *et al*. 2023), thus provides a comparable context for evolutionary separation between *N. vectensis* and *S. callimorphus* for genome content as well as organization.

To improve our understanding of the *Edwardsia* genus and increase the genomic resources for actinarians, we have sequenced and assembled a *de-novo* genome for the Edwardsiid anemone *Edwardsia elegans* Verrill 1869. *E. elegans* live in soft bottom habitats from the intertidal to 120m depth along the North Atlantic coast of North America from Maine to North Carolina. The latitudinal range of *E. elegans* overlaps with other Edwardsiidae species, most notably the model cnidarian *N. vectensis*. The *E. elegans* range also coincides with some of the fastest warming waters on Earth, where genome-enabled species can be utilized to monitor acclimation and adaptation to climate change (Kavanaugh *et al*. 2017; Aguirre-Liguori *et al*. 2021).

Here we used this genome to compare genomic variation between sea anemones, including two Edwardsiid species, to compare patterns of representation for gene families, specific conserved features (microRNAs, Ultra Conserved Noncoding Elements (UCNEs)), and syntenic organization. Syntenic regions have developed into particularly useful regions of the genome to highlight evolutionary relationships (Schultz *et al*. 2023) in addition to their influence in gene regulation (Marques-Bonet *et al*. 2004; Farre *et al*. 2019). Altogether we show that the addition of this genome from *Edwardsia elegans* provided an insightful comparison to better understand variation in genome content and organization for actiniarians including model species like *Nematostella vectensis*.

## Methods

### Specimen Collection and Culture

Adult *E. elegans* were collected with the assistance of Gulf of Maine, Incorporated (www.gulfofme.com) from Cobscook Bay, Maine. They were shipped to the University of North Carolina at Charlotte, where they were housed in a recirculating water system and in finger bowls. The recirculating water system was set to a temperature of 16 ºC with artificial seawater (Instant Ocean) at 30 parts per thousand (ppt), with a ∼5% water change twice per week. *E. elegans* in finger bowls were kept in 30 ppt artificial seawater at room temperature with a full water change once a week. All anemones were cultured with sand substrate (Nature’s Ocean Marine White #0) and regularly fed freshly hatched *Artemia* nauplii and pieces of mantle tissue from mussel (*Mytilus edulis*). Prior to DNA extraction, anemones from the recirculating tanks were placed in standing glass finger bowls with a small amount of sand substrate and held in a 16 ºC incubator in total darkness and starved for two weeks to minimize DNA from food sources.

### DNA Extraction, Library Preparation and Sequencing

A single individual *E. elegans* was used for DNA extraction for both Illumina and Oxford Nanopore sequencing. DNA was extracted using a protocol based on Smith et al (Smith *et al*. 2023), quantified using a Qubit (Q32857) with the dsDNA High Sensitivity kit (Q32851), following standard manufacturer protocols, quality checked on a 1% agarose gel.

A library for Illumina sequencing was prepared with the Illumina DNA PCR-Free Prep kit (20041855). The library was quantified and quality checked using a Qubit (Q32857) and dsDNA Quantitation, High Sensitivity kit (Q32851). The resulting library was sequenced using NextSeq 2000 P3 (2×150bp) on a NextSeq 2000 instrument at the University of North Carolina at Charlotte.

Four libraries were prepared for Nanopore sequencing using the ligation sequencing kit (LSK109; Oxford Nanopore Technologies) and were sequenced with a MinION sequencer (R10.4; Oxford Nanopore Technologies). All libraries were sequenced on a single flow cell for approximately 45 hours each, with the flow cell being washed between each new library following manufacturer protocols. Prior to assembly, raw nanopore reads were basecalled using Guppy v. 6.0.6 (Oxford Nanopore Technologies) with Cuda and a configuration file for the R10.3 flow cell and a minimum quality score of 7.

### Assembly

Prior to genome assembly, Illumina reads were used to estimate the genome size. Jellyfish v. 2.3.0 (Marcais AND Kingsford 2011) was used to generate counts for 21 bp kmers and were visualized in GenomeScope (Vurture *et al*. 2017), estimating the genome to be roughly 332Mb with heterozygosity of 2.5%. Based on the high heterozygosity from the kmer estimates, we used the program wtdbg2 v. 2.5 (Ruan AND Li 2020), for genome assembly. All settings were default except ‘--edge-min’ which was reduced to 2, sampling rate ‘-S’ which was reduced to 2, and ‘--rescue-low-cov-edges’ was added. After assembly, the genome was polished twice using Illumina raw reads with the program Pilon v. 1.24 with default settings (Walker *et al*. 2014).

Repeat content was determined using a combination of RepeatModeler v 2.0.2 (Smit and Hubley 2008-2015) and RepeatMasker v. 4.1.2 (Smit *et al*. 2013-2015). First the assembled genome was run through RepeatModeler using default settings. This generated a masked genome that was used as an input for RepeatMasker, with default settings. The masked genome output from RepeatMasker was used for protein predictions.

### Annotations

Genes in the final assembled genome were annotated using the program BRAKER2 v 2.1.5 (Bruna *et al*. 2021) which uses GeneMark and Augustus to predict gene models. Three data sources were used as inputs into BRAKER2: 1) raw RNA-seq data from *E. elegans* (GenBank Accession #GKWK00000000), 2) proteins from the confamilial species *N. vectensis* downloaded from SIMRbase (NV2, https://simrbase.stowers.org/starletseaanemone) and 3) the metazoan protein database available from ProtHint (https://bioinf.uni-greifswald.de/bioinf/partitioned_odb11/). Protein files from *N. vectensis* and the ProtHint metazoan database were concatenated into a single file. Outputs from BRAKER2 were then queried with BLASTp v2.11.0+ against proteins in databases from NCBI (accessed March 2022), Uniprot Swiss-Prot (accessed March 2022), and *N. vectensis* proteins using Diamond BLAST (Buchfink *et al*. 2014) to identify known genes for annotations. BLASTp matches were sorted by e-value and percent similarity, and top hits based on e-value and bit score were utilized for each predicted gene. Gene predictions were also annotated with functional domains for protein families using the program HmmrScan v 3.1 (Eddy 2009). Only hits with an E-value less than 1e-05 were retained and used for annotation.

BUSCO v. 5.1.3 (Manni *et al*. 2021) using the metazoa_odb10 ortholog set was used to assess quality based on completeness of both gene predictions and the genome as a whole.

### Orthologous Proteins

Orthofinder v. 2.4.0 (Emms AND Kelly 2019) was used to identify orthogroups, orthologs, and gene duplications between the newly assembled *E. elegans* genome and other anthozoans. Single copy orthologs generated were then used for syntenic analysis (see below). We used proteins from the following anthozoans: *Exaiptasia diaphana, Actinia tenebrosa, N. vectensis, S. callimorphus*, and *E. elegans*. The hexacoral *Acropora millepora* and the octocoral *Renilla reniformis* were used as outgroups. *E. elegans* proteins were from this study, *N. vectensis* and *S. callimorphus* proteins were obtained from SIMRbase, and all other proteins were obtained from NCBI. Orthofinder was run with default settings.

### Syntenic Analysis

We compared both macro- and micro-syntenic regions between our *E. elegans* genome and the other Edwardsiidae anemones species with high quality genome assemblies, *N. vectensis* and *S. callimorphus*. For macrosyntenic analysis, reciprocal protein BLASTp were performed between *E elegans* protein predictions generated from BRAKER2 and *N. vectensis* protein predictions, as well as *E. elegans* and *S. callimorphus*. For Oxford Dot Plot comparisons, top hits for reciprocal BLASTp were taken and one-to-one hits between species were identified. Information on locations for each of these proteins was then taken. Genome location information for each gene was compared with the R (R CORE TEAM 2022) program MacrosyntR (El HILALI AND COPLEY 2023) to generate Oxford Dot Plot comparisons for pairwise species comparisons. For ribbon plot comparisons, orthogroups with only a single copy in all three Edwardsiidae anemones were identified and confirmed with reciprocal BLASTp searches. MacrosyntR was then used to generate ribbon plots.

For microsyntenic analysis, sequences for each region of interest were obtained and used for BLASTp searches against proteins annotated from BRAKER2 and then the locations of the top hits were found in the genomes of each target species. Top hits for each gene were taken and then sorted by starting location along each chromosome/scaffold, then gene order, proximity, and direction were compared between each species.

### Ultra Conserved Noncoding Elements

To identify ultra conserved noncoding elements (UCNE) in the *E. elegans* genome, known UCNEs annotated for *N. vectensis* and *S. callimorphus* were obtained from SIMRbase (https://simrbase.stowers.org/starletseaanemone). 143 UCNE were then searched for in the *E. elegans* genome with BLASTn. Hits higher than an e-value of 1e-05 and shorter than a length of 100bp were removed, as all but two of the UCNEs identified in *N. vectensis* are a minimum of 100bp. Top hits from the *E. elegans* genome were annotated and used to generate an Oxford dot plot in MacrosyntR with UCNEs from *N. vectensis* and *S. callimorphus*.

### Small-RNA sequencing and annotation of miRNAs

Total RNA was extracted from *E. elegans* polyps using TRIzol reagent (Thermo Fisher Scientific, USA), following the manufacturer’s protocol. The extracted RNA was then selected for 18-30 nucleotides on a 15% denaturing urea polyacrylamide gel (Bio-Rad, USA). RNA was eluted overnight in 0.3 M NaCl. For library preparation, we used a modified version of the Illumina TruSeq small-RNA Cloning Protocol (Zamore lab, http://www.umassmed.edu/zamore/resources/protocols/, accessed April 2014). In brief, the small RNAs (sRNAs) were ligated to 3’ and 5’ adapters containing four random nucleotides at the ligation interface to minimize ligation bias. The ligation products were then reverse transcribed using SuperScript III Reverse Transcriptase (Thermo Fisher Scientific). The cDNA samples were PCR amplified using the KAPA Real-Time Library Amplification Kit (PeqLab, Germany). The amplified cDNA was purified on 2% low-melt agarose gels (Bio-Rad). The small RNA library was validated on a High Sensitivity D1000 ScreenTape (Agilent, USA) Finally, the sRNA library was sequenced on NextSeq 500 (Illumina, USA), with read lengths of 50 nucleotides. miRNA analysis was performed using mirDeep2 (FRIEDLÄNDER *et al*. 2011). Initially, the sequencing data was pre-processed to remove adapters and the short sequences less than 18 nucleotides were discarded. The miRDeep2 core algorithm was then used to identify miRNAs, with the sequenced *E. elegans* genome from the current study used as a reference. The identified miRNA candidates were validated manually based on specific criteria suggested for miRNA annotation in animals (Fromm *et al*. 2015), including a 2-nucleotide overhang on the 3’ end of precursor miRNA, a distinct length of ∼18-23 nucleotides, predicted folding of pre-miRNA transcript into a hairpin structure of ∼60 nucleotides, a clear signature for strand selection with a dominant guide strand, that contains a homogeneous 5’ end, guide/star ratio higher than two, and at least 16 nucleotide complementarities between mature and star strand. However, the terminal loop size of precursor miRNAs above eight nucleotides and the consistency of the star strand of miRNAs were not considered since cnidarian miRNAs do not seem to follow these rules (Praher *et al*. 2021). To identify conserved miRNA sequences across cnidarians, mature miRNA and miRNA precursors from ten cnidarian species were retrieved from previous miRNA studies (Liew *et al*. 2014; Moran *et al*. 2014; Gajigan AND Conaco 2017; Baumgarten *et al*. 2018; Fridrich *et al*. 2020; Praher *et al*. 2021) and used as input in the miRDeep2 quantification algorithm. Output files from the miRDeep2 core algorithm and accepted miRNAs are provided in the supplementary table.

## Results

### Genome Assembly and Annotation

Nanopore sequencing of genomic DNA generated approximately 9.5 gigabytes of raw data. Raw nanopore reads were basecalled leading to 385,657 reads that were then assembled into 6,319 contigs. Illumina sequencing of genomic DNA resulted in 79,640,638 paired end reads and were used to polish the 6,319 assembled contigs. This results in an *E. elegans* genome that is 396,821,203bps in length with an N50 of 151,884bp. Genome quality was assessed with BUSCO and resulted in a score of 88% (Single:87.6%, Duplicate:0.7%, Missing: 5.0%) based on the metazoan database. GC content is 39.9% and repeat content was calculated to be 47.8% of the genome (Figure 1). Beside unclassified repeats, the most abundant type of repetitive elements is the LINE group, making up 3.78% of the genome. LTRs make up 1.09% of the genome and DNA transposons make up 1.49% of the genome. 49,837 protein coding regions were predicted using Braker2 (Bruna *et al*. 2021).

**Figure 1:**
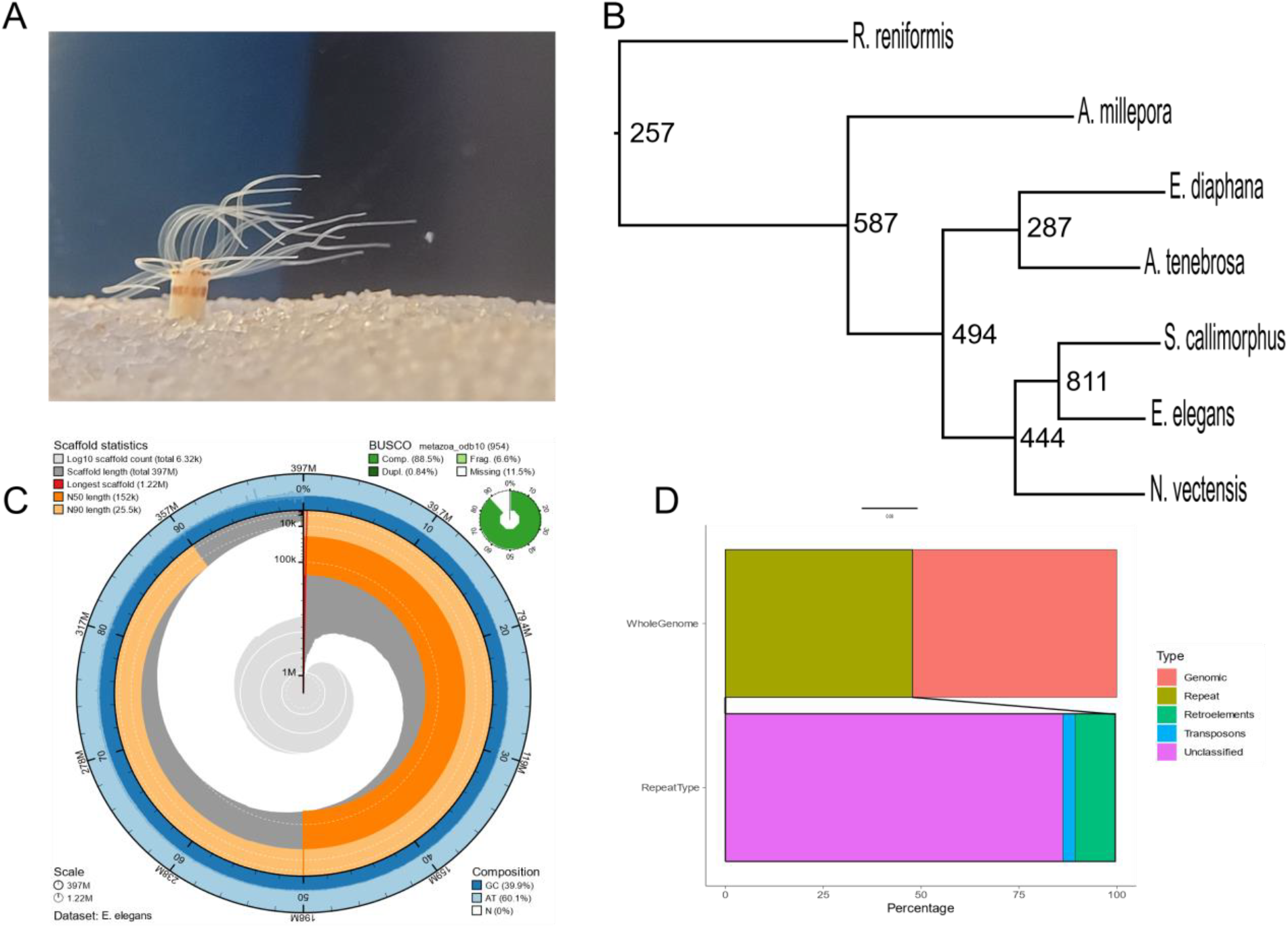
A) *Edwardsia elegans* in the recirculating aquaria at UNC Charlotte. B) Phylogenetic tree with node number representing duplications. C) Snail plot of the *E. elegans* genome generated using BlobToolKit (Challis *et al*. 2020). D) Repetitive Regions of the genome.

### Orthologous Protein Groups

Orthofinder (Emms AND Kelly 2019) was used to identify and compare orthologous groups shared between *E. elegans* and other sea anemone species. Orthofinder generated 24,525 orthogroups, with 91.2% of total proteins from all species being placed into an orthogroup. There were 6,134 orthogroups (25%) containing proteins from all species, with 869 single copy orthogroups shared between all species. All species had over 90% of their proteins placed into an orthogroup, except *E. elegans* and *R. reniformis*, which had 82.0% and 83.5% respectively. 74.7% of all orthogroups contained at least one *E. elegans* protein, the highest of any species examined here. We determined the number of gene duplications at each node (Figure 1b). Focusing on the Edwardsiidae branch, there are 444 duplications for the node leading to the Edwardsiid species and nearly double that number for the *E. elegans/S. callimorphus* node.

When comparing the number of single copy orthologs in species in the Edwardsiidae family (i.e., *E. elegans, N. vectensis*, and *S. callimorphus*), we find 3,068 single copy orthologs shared between all three Edwardsiidae species (Figure 2). When comparing *Edwardsia elegans* to the other Edwardsiidae species, 5,569 single copy orthologs are shared with *N. vectensis*, or approximately 11.7% of the predicted proteins in *E. elegans*. There are 5,420 single copy orthologs shared with *S. callimorphus*, approximately 10.8% of the predicted proteins in *E. elegans*. Single copy orthologs shared between all three Edwardsiidae anemones include several housekeeping genes such as heat shock proteins, GMP synthase, *wnt*, and glyceraldehyde-3-phosphate dehydrogenase. There were 2501 single copy orthologs shared between *E. elegans* and *N. vectensis* but not *S. callimorphus*, and 2352 single copy orthologs shared by *E. elegans* and *S. callimorphus* but not *N. vectensis*. The single copy orthologs shared in all three species were then used to identify regions of synteny between *E. elegans, S. callimorphus*, and *N. vectensis*.

**Figure 2:**
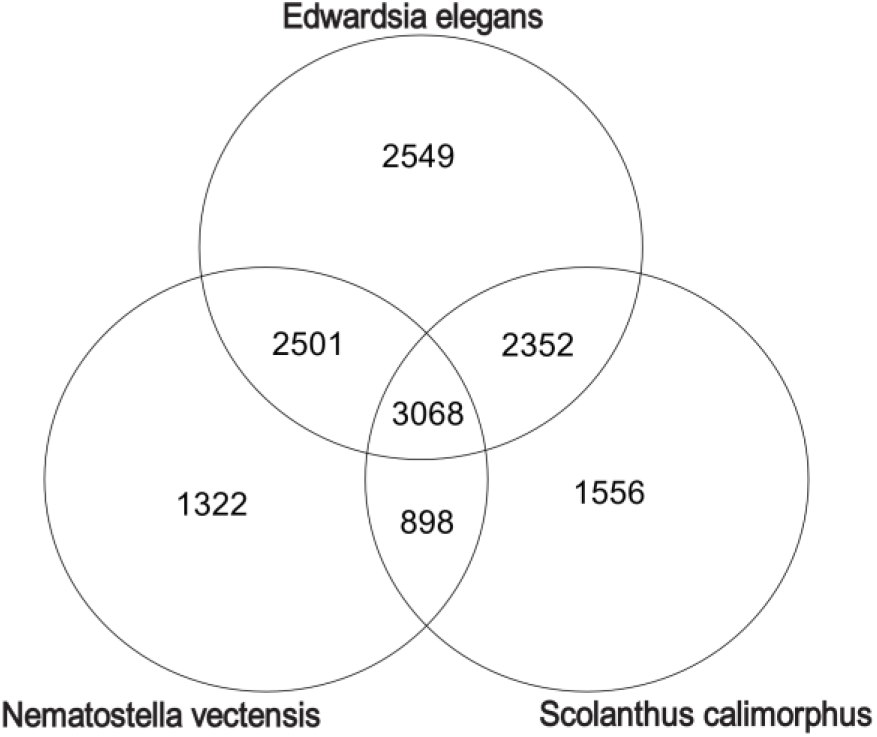
Venn Diagram showing the number of species specific and shared Single Copy Orthologs found between the three Edwardsiid species.

### Syntenic Regions of Genomes

#### Macrosyntenic Arrangements

We identified numerous conserved macrosyntenic regions between *E. elegans* and the two other Edwardsiidae anemones, *N. vectensis* and *S. callimorphus*. Using reciprocal BLASTp across the largest 25 contigs in the *E. elegans* genome there were 1024 one-to-one orthologs between *E. elegans* and *N. vectensis* and 789 one-to-one orthologs between *E. elegans* and *S. callimorphus*. These comparisons show many *E. elegans* contigs localizing to single chromosomes in *S. callimorphus* and *N. vectensis* (Figure 3A & B, respectively). These macrosyntenic regions shared between *E. elegans* and *S. callimorphus* are more tightly distributed, as seen in more instances of straight lines in Figure 2B. For example, *E. elegans* ctg_0001 has two syntenic regions of over 10 single copy orthologs that are also located in the same order on chromosome 4 of *S. callimorphus*. However, these same proteins are more dispersed on chromosome 2 of *N. vectensis*, with only one conserved syntenic region of five proteins. We also show that *E. elegans* ctg_0023 has a syntenic region of 21 single copy orthologs shared on chromosome 4 of *S. callimorphus*, while the same region on chromosome 2 of *N. vectensis* has only a small syntenic region of 5 proteins while the rest are mixed across the chromosome.

**Figure 3:**
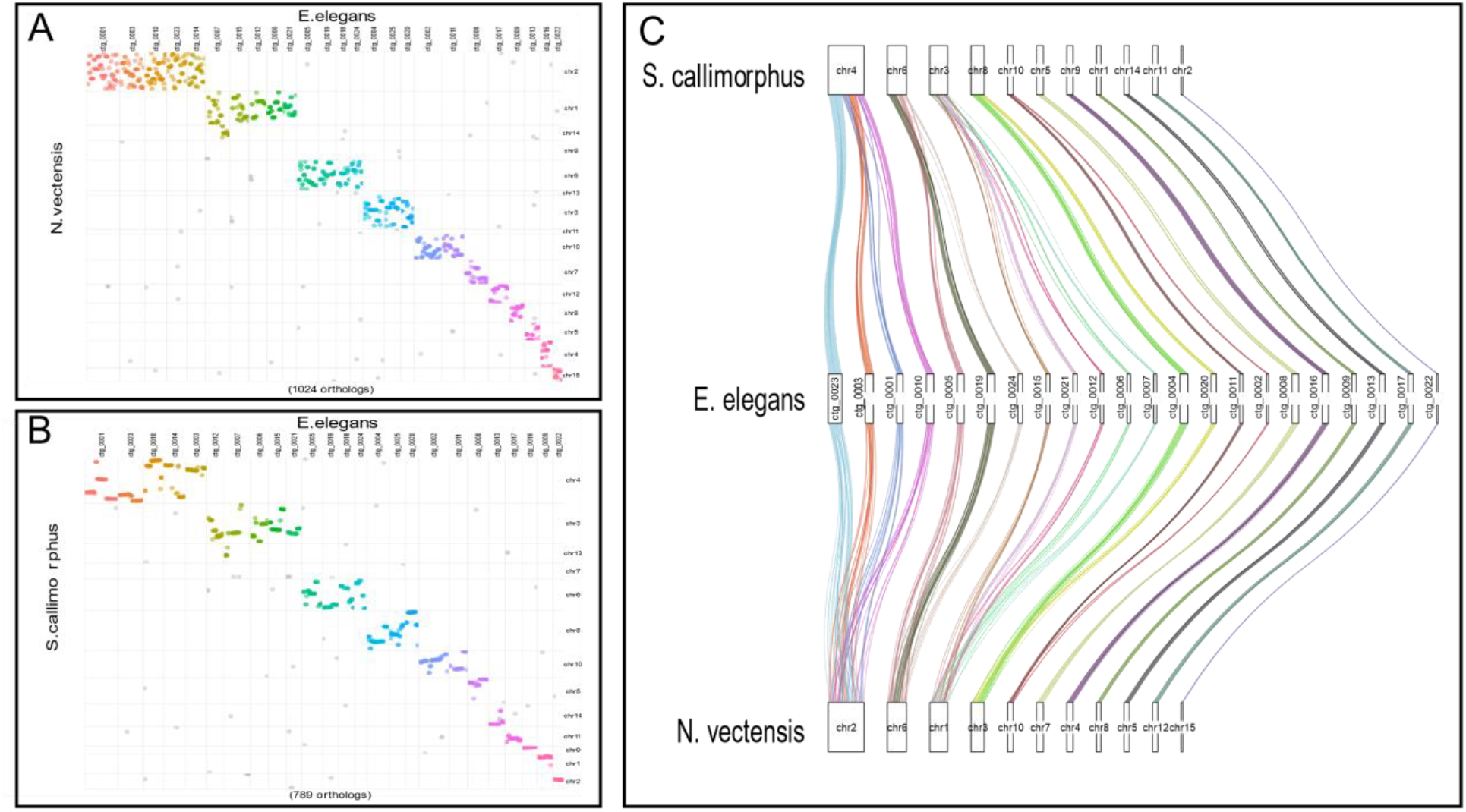
Macrosyntenic regions between *E. elegans* and a) *N. vectensis* and b) *S. callimorphus*. Each dot represents the best one-to-one reciprocal BLASTp hit for a protein between the two species. C) A ribbon plot of single copy orthologs showing macrosynteny between *E. elegans* and *S. callimorphus* and *N. vectensis*.

We identified regions where *E. elegans* and *S. callimorphus* share longer stretches of genes in the same order that are not in synteny with *N. vectensis*. In these largest 25 contigs of the *E. elegans* genome, there are 13 regions of 10 or more genes in nearly identical gene order between *E. elegans* and *S. callimorphus*, where these same regions in *N. vectensis* are dispersed and are not larger than 9 protein coding genes. The largest syntenic region between *E. elegans* and *S. callimorphus* in this analysis is 25 protein coding genes found across approximately 450kb in both species, whereas the largest syntenic region between *E. elegans* and *N. vectensis* is 9 coding genes across approximately 100kb.

Using single copy orthologs shared between all three Edwardsiidae species, we investigated how macrosyntenic regions have been rearranged (Figure 3C). For example, syntenic blocks of genes located on *E. elegans* ctg_0003 and ctg_0010 correspond to two regions of chromosome 4 in *S. callimorphus*. However, these clustered genes are much more diffuse in *N. vectensis*. Similarly, two syntenic regions on *E. elegans* ctg_0004 and ctg_0020 match chromosome 8 of *S. callimorphus*, while these gene regions are dispersed on chromosome 3 of *N. vectensis*. These regions are indication that orthologous genes maintain intrachromosomal locations between these Edwardsiid species, but the syntenic organization is less conserved in *N. vectensis*.

#### Microsyntenic Trends

We compared the microsynteny of previously identified developmental genes that are clustered in Edwardsiidae anemones with the model anemone *Exaiptasia diaphana (Baumgarten et al. 2015)* as an outgroup. The first region we compared was a cluster of three Paired-class homeobox genes, Hbn-Rx-Otp, involved in animal development (Holland 2013) and previously shown to be clustered in *N. vectensis* (Mazza et al. 2010). All three genes are found in close association in all four of these anemone species, however the genomic organization varies (Figure 4A). *E. elegans* and *S. callimorphus* share the same order of Rx-Otp-Hbn and all genes are in the same transcriptional orientation. The gene order of these two species differs compared to *N. vectensis* and *E. diaphana*, where the orientation and transcriptional direction of the Otp and Rx genes is inverted with respect to Hbn.

**Figure 4.**
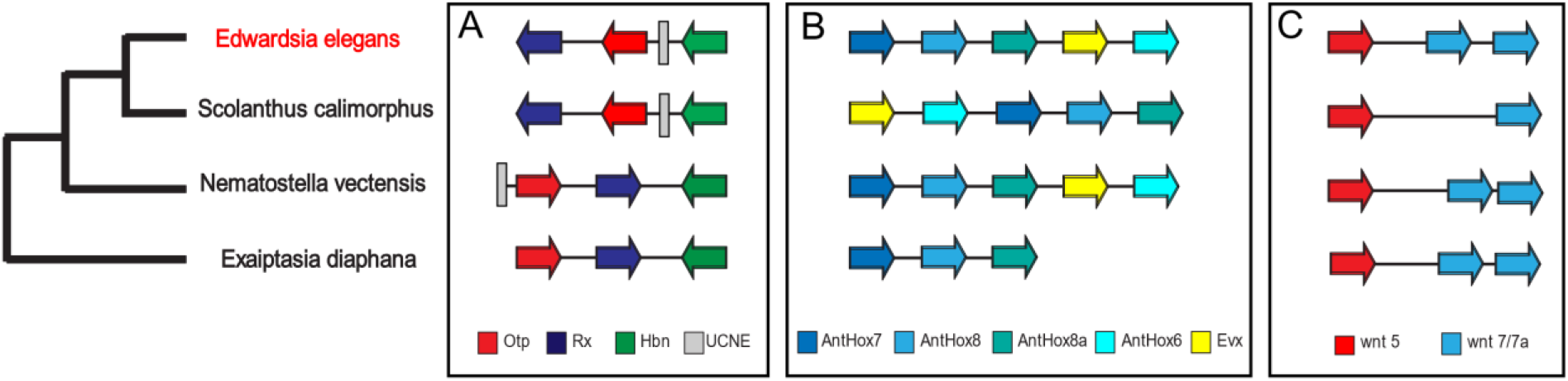
: Microsyntenic comparisons between *E. elegans* and the three other actinarians. A) Hbn-Otp-Rx and an UCNE B) HOX C) wnt5 and 7/7a

Next, we compared regions of HOX genes, which are critical for animal embryonic development and whose gene order impacts axial patterning (Gellon AND Mcginnis 1998; Dubuc *et al*. 2018). We identified 30 HOX genes previously annotated in *N. vectensis* and other sea anemones (Chourrout *et al*. 2006; Ryan *et al*. 2007; Zimmermann *et al*. 2023). In *E. elegans* we find four clusters of three or more Hox genes along with three cluster or two clusters. These clusters were not all located on the same contig as they are in other sea anemones, which may be an artifact of the fragmented assembly compared to the chromosome scale of the other anemones’ genomes. The ParaHox cluster (x*lox/cdx & gsx*) is conserved in all four species with highly similar distances between genes. A cluster of hox genes which contains *anthox7*, a*nthox8, anthox8b, evx*, and *anthox6*, is identical between *E. elegans* and *N. vectensis*, while this cluster is slightly rearranged in *S. callimorphus*, with *evx* and a*nthox6* in a different position relative to the other genes (Figure 4B). We also annotated a pair of *nk* genes, *nk1* and *nk5*, that form a paired cluster in *E. elegans* on ctg_1036, as well as a cluster that contains *msx, nk2c*, and *nk2d* on ctg_0003. *hlxe* and *gbx* are arranged as a pair on ctg_0208. We also identified a cluster of *anthox1, lbx, nk3, rough* on ctg_0005.

*E. elegans* also has a *wnt* cluster that was identified in *N. vectensis* and other cnidarians (Ryan *et al*. 2006; Ryan *et al*. 2007; Sullivan *et al*. 2007; Steinworth *et al*. 2023). Both *E. elegans* and *S. callimorphus* have a cluster of *wnt-5* and *wnt-7/7b*. Our protein predictions are not capable of recognizing the alternative splicing of *wnt-7/7b* found in *N. vectensis*, but we do get two distinct proteins are grouped into the same orthogroup from Orthofinder, which also contains other *wnt7b* isoforms from *N. vectensis*. In this same orthogroup we got only a single *S. callimorphus wnt7b* ortholog was present. This *wnt* cluster spans a similar size of the genome in both *N. vectensis* and *E. elegans* while in *S. callimorphus* these genes are distributed over approximately 12,000 additional bases.

### Ultra Conserved Noncoding Elements

Previous research with two species has shown ultra conserved noncoding elements (UCNEs) present in the family Edwardsiidae (Zimmermann *et al*. 2023). Here we show there are at least 92 UCNEs (of 143 total) that have previously been found in *N. vectensis* and *S. callimorphus* are present in this new *E. elegans* genome. These UCNEs are distributed across 62 different contigs in the genome. We also observe UCNEs found near syntenic regions, such as UCNE 579 within the Otp-Rx-Hbn syntenic region, approximately 100bp upstream from OTP. This UCNE is found outside of this cluster in *N. vectensis* but is present inside the cluster in both *E. elegans* and *S. callimorphus* (Figure 4A), presumably due to the same inversion event.

We further observed that there are several regions where these UCNE are found in syntenic arrangements in all three Edwardsiid species. These regions are also found to be nearly uniformly distributed across many regions between all three species (Figure 5). Similar to the macrosynteny for genes in orthogroups, these UCNE regions are more contiguous between *E. elegans* and *S. callimorphus* (Figure 5A), with more instances of rearrangement between *E. elegans* and *N. vectensis* (Figure 5B).

**Figure 5:**
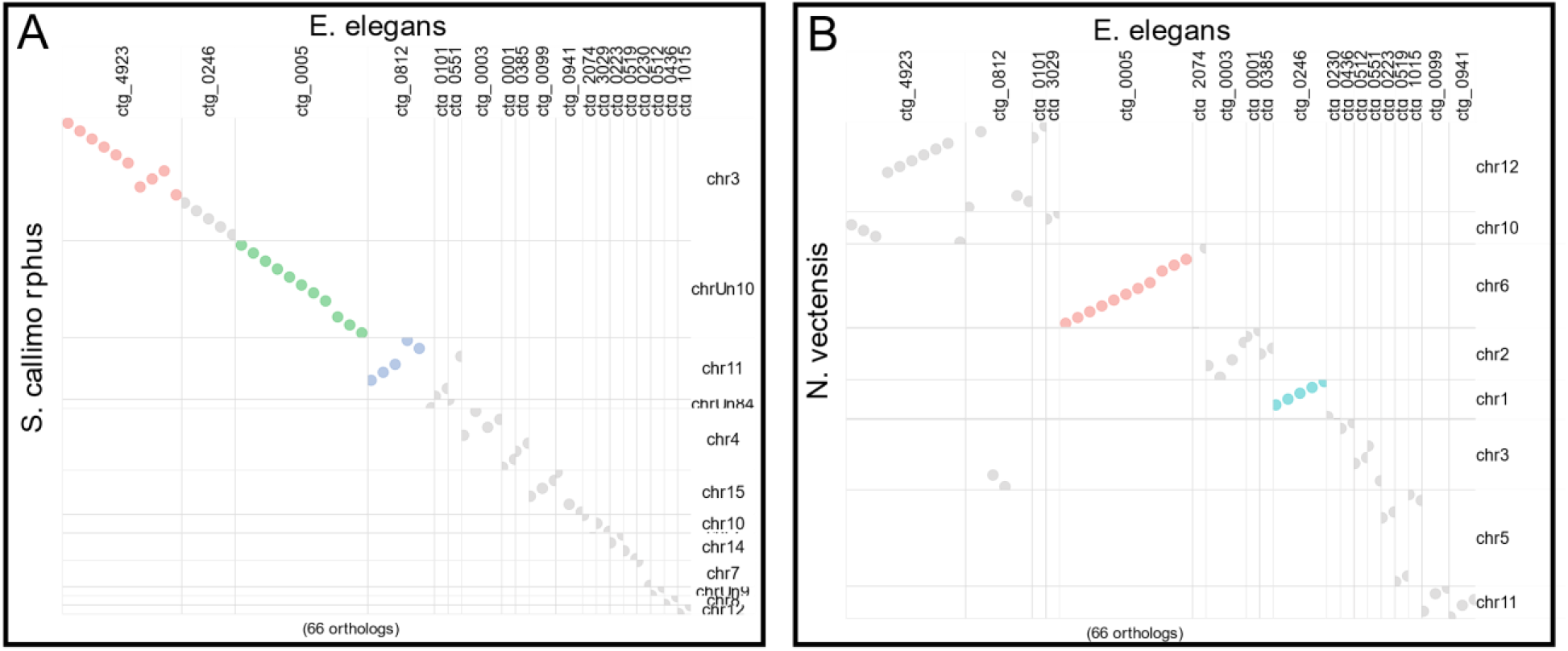
Dot plot of UCNE regions between A) *E. elegans* and *S. callimorphus* and B) *E. elegans* and *N. vectensis*

### *E. elegans* miRNA repertoire

To explore miRNA repertoire of *E. elegans*, we prepared small RNA libraries from the polyps and analyzed the data using miRDeep2 (FriedlÄnder *et al*. 2011) (see Material and methods for details). The length distribution of total small RNA reads revealed two major small RNA populations with nucleotide lengths of 19-22 and 26-31 nucleotides (Figure 6A). The first population represents putative miRNAs and small-interfering RNAs (siRNAs), whereas the second population represents putative PIWI-interacting RNAs (piRNAs) and constituted the majority of the sequenced small RNAs. The length distribution of the small RNAs and the enrichment of piRNA population over miRNAs is consistent with previous studies from other cnidarians (Liew *et al*. 2014; Moran *et al*. 2014; Gajigan AND Conaco 2017; Baumgarten *et al*. 2018; Fridrich *et al*. 2020; Praher *et al*. 2021). The miRDeep2 predicted 137 miRNA candidates, of which 30 were identified as bona fide miRNAs based on specific criteria suggested for miRNA annotation in animals (see Material and methods for details). The analysis of the nucleotide composition of miRNA sequences identified here reveals a strong bias for U at the first position of the mature sequence, representing a characteristic feature of miRNAs (Figure 6B). This bias is explained by the preference of Argonaute proteins for a U at the 5’ end of the miRNA, which is a known characteristic of miRNAs in bilaterian (Frank *et al*. 2010) and cnidarians (Moran *et al*. 2014; Praher *et al*. 2021).A comprehensive list of bona fide miRNAs of *E. elegans* and those who did not pass the criteria is available in supplementary Table X. Using miRDeep2’s quantifier module, we also detected conserved miRNAs across cnidarians by providing sequences of previously identified cnidarian miRNAs (Liew *et al*. 2014; Moran *et al*. 2014; Gajigan AND Conaco 2017; Baumgarten *et al*. 2018; Fridrich *et al*. 2020; Praher *et al*. 2021). Only ten out of the 30 miRNAs were homologous in sequence to ones that had been annotated previously in one or more anthozoan species, with only seven being conserved among all the anthozoans for which there are data available (Figure 6C).

**Figure 6:**
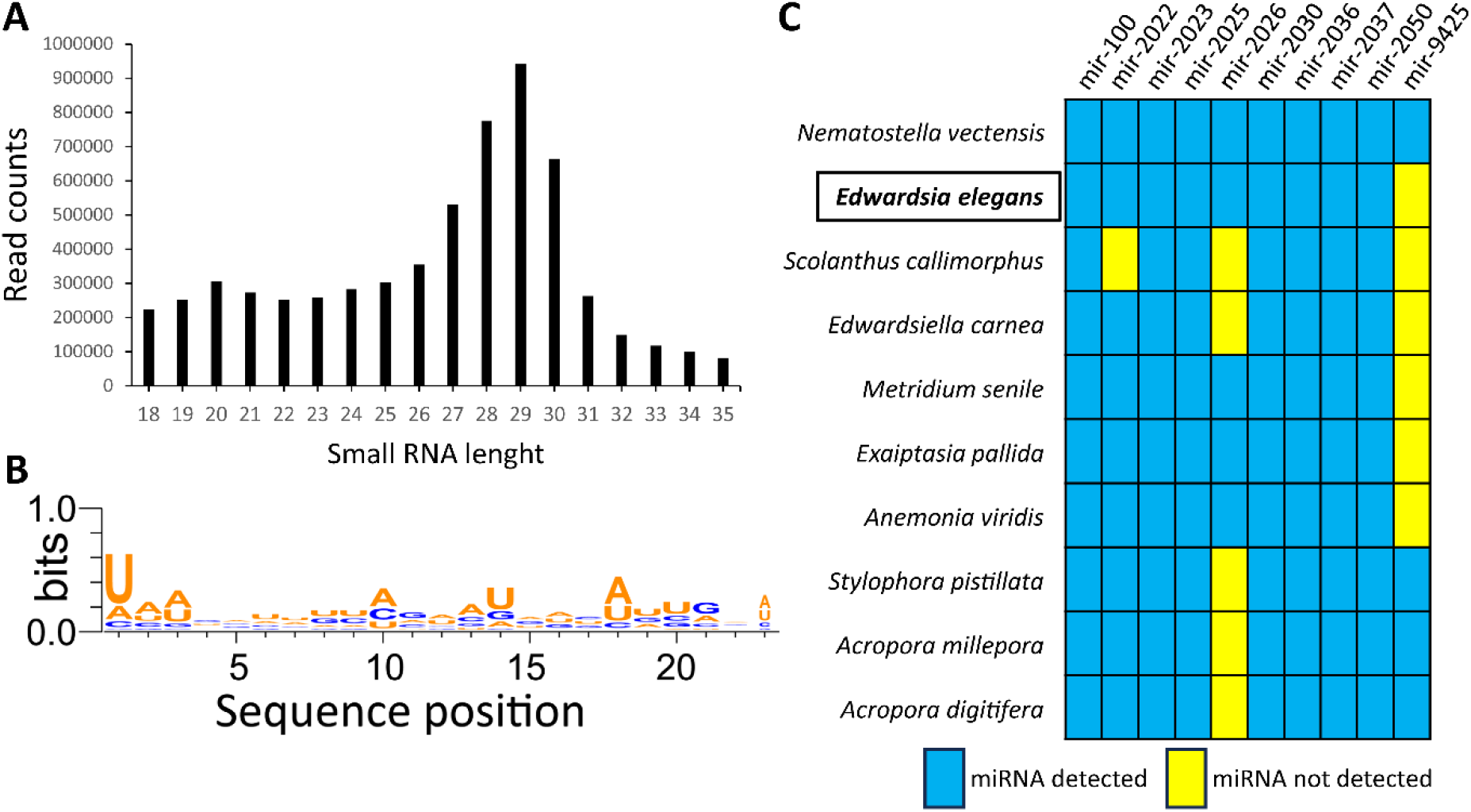
Annotation of the *E. elegans* miRNA repertoire. A. Two distinct populations of small RNA reads: putative siRNAs/miRNAs (19-23nt) and putative piRNAs (27-31nt). B. miRNA sequences exhibit a bias toward uridine at position one. WebLogo3 was utilized to create sequence logos (Crooks *et al*. 2004). **C**. The miRNAs that are evolutionarily conserved between *E. elegans* and other sequenced anthozoan species.

## Discussion

The de-novo genome for the sea anemone *Edwardsia elegans* is the first genome for a species in the *Edwardsia* genus. The genome is 396.8Mb in length and predicted to contain 49,837 protein coding regions, with a repeat content calculated to be approximately 47%. The three Edwardsiidae sea anemones we compared share over 3,000 single copy orthologs with each other, making up greater than 6% of each of their proteomes. The true number is likely higher as the annotations for *N. vectensis* include isoforms of multiple genes, such as a few *wnt*, where they are single copies in *E. elegans* and *S. callimorphus* but annotated as two isoforms for *N. vectensis*.

Comparative analysis of all identified orthogroups revealed *E. elegans* being most closely related to *S. callimorphus*, with *N. vectensis* as more distantly related. This pattern of orthogroups is consistent with previous research on the family Edwardsiidae and strengthens our understanding of the relationship of Edwardsiid sea anemones (Daly 2002; Rodriguez *et al*. 2014; Zimmermann *et al*. 2023). While the number of proteins predicted are higher than the other Edwardsiidae anemones (Zimmermann *et al*. 2023), high BUSCO scores and the fact that the majority of predicted proteins had significant BLAST hits give us confidence that the are transcribed portions of the genome. Efforts to reduce the number of genes with tools in the BRAKER pipeline resulted in removal of genes previously annotated in *E. elegans* (e.g., toxin genes). Thus, we preferred to retain all predicted genes for transparency in the annotation process for this *de novo* genome. While the true number of proteins in the *E. elegans* genome is likely smaller than what we have predicted, we are leaving all predicted proteins rather than risk removing true proteins.

Macrosynteny analyses have recently emerged as a particularly informative approach for inferring evolutionary relationships (Simakov *et al*. 2020; Schultz *et al*. 2023). We observe macrosyntenic regions of both proteins and UCNEs that are shared between all three Edwardsiidae species, but many of these regions are more contiguous and linear between *E. elegans* and *S. callimorphus*. We expect this in more closely related species that have had less evolutionary time to differentiate. Sequencing of other Edwardsiidae anemones, especially those in the *Edwardsia* genus, will be needed to determine if these regions are conserved more broadly in the genus. More genomic detail will also allow us to see how these regions differentiated in the family Edwardsiidae to discern if rearrangements in *N. vectensis* are unique in the Edwardsiidae family.

Gene clusters of evolutionary conserved transcription factors (e.g., Hox genes) are hypothesized to be conserved over large phylogenetic distances due to functional constraints. The spatial arrangement of these genes can be related to the timing and location of expression during development (Gellon AND Mcginnis 1998). Cnidarians, particularly *N. vectensis*, have been insightful for the evolutionary history of the *Hox, wnt*, Paired-Class, and other genes due to the arrangement of these genes in the genome and their spatiotemporal expression patterns and their developmental functions (Kusserow *et al*. 2005; Ryan *et al*. 2006; Ryan *et al*. 2007; Sullivan *et al*. 2007; Dubuc *et al*. 2018; Steinworth *et al*. 2023). Comparisons of *E. elegans* with other Edwardsiid anemones suggests these clusters vary within this family of cnidarians. Previous research in *N. vectensis* have shown these gene clusters to be expressed at particular times and locations in development (Ryan *et al*. 2007; Sullivan *et al*. 2007; Mazza *et al*. 2010; Kraus *et al*. 2016; Dubuc *et al*. 2018; He *et al*. 2018; Holstein 2022), and thus research into the developmental timing using *E. elegans* or *S. callimorphus* in comparison with *N. vectensis* would be needed to determine if the order of these genes influences their developmental timing in these Edwardsiid species and possibly in other actinarians.

miRNAs are small non-coding RNAs that play a crucial role in various biological processes such as development and cell physiology in both plants and animals (Bartel 2009; Voinnet 2009; Modepalli *et al*. 2018). Despite extensive research, the evolution of miRNAs remains enigmatic, particularly with respect to whether they have a common origin in plants and animals due to their differences in biogenesis (Axtell *et al*. 2011; Moran *et al*. 2013; Moran *et al*. 2017; Tripathi *et al*. 2022; Edelbroek *et al*. 2024). The phylogenetic position of cnidarians offers a unique opportunity to explore miRNA evolution. Here, we analyzed the miRNAs in *E. elegans* and identified 30 bonified miRNAs. By comparing miRNA sequences with other cnidarian species, we discovered nine miRNAs that were conserved with *N. vectensis*, a closely related species, and six that were shared with other anthozoan species. The findings not only revealed *E. elegans*-specific miRNAs but also supported our earlier report on miRNA sequence evolution in cnidarians (Praher *et al*. 2021), which highlighted rapid gains and losses of miRNAs in Cnidaria, indicating a higher miRNA turnover rate in cnidarians compared to bilaterians.

We have assembled and annotated the genome of *Edwardsia elegans*, which will provide a valuable tool for studying cnidarian genomic diversity and evolution. More genetic information for species within the actiniarians will improve our understanding of the evolutionary history of both actinarians and cnidarians as a whole and can shed light on many processes that are shared between the two. Broad comparisons across taxonomic groups will benefit by increasing the available species that can be used for comparisons. Increasing the number of species with genomic resources is the first step and will also improve our understanding of genomic diversity and variation across a lineage. This genome will help aid in comparisons of the diversity of ecologically and phylogenetically important cnidarians, as well as comparative investigation of the parallel evolution of cnidarians and bilaterians.

## Data availability

The genome and raw data sequences has been deposited on NCBI, accession number JBHDXZ000000000. Codes for this study are available at github.com/austonrut/EelegansGenome. Genomes and proteins that were not generated as part of this study we accessed from NCBI.

## Funding

This research was supported through incentive funds from the University of North Carolina at Charlotte. Small RNA sequencing was supported by Binational Science Foundation program with the National Science Foundation grants 2020669 and 1536530 to Y.M. and A.M.R.

## Conflict of Interest

The authors declare no conflict of interest.

## Acknowledgements

CIPHER Center for support with reagents and sequencing resources. UNC Charlotte High Performance Computing for technical support and access to computing resources. A huge thank you to Dr. Edward Smith for advice and assistance with both high molecular weight DNA extractions and bioinformatics. Thank you to Dr. Sydney Birch for help with bioinformatics.

## Author Contributions

A.I.R. and A.M.R conceived of study. A.I.R. and R.K. generated sequence data. A.I.R. analyzed all data except miRNA, which was generated and analyzed by V.M. and Y.M. A.I.R., A.M.R., V.M., and Y.M wrote the manuscript. All authors edited the manuscript.

